# Correlational selection and genetic architecture promote the leaf economics spectrum in a perennial grass

**DOI:** 10.1101/2021.11.14.468541

**Authors:** Robert W. Heckman, Jason E. Bonnette, Brandon E. Campitelli, Philip A. Fay, Thomas E. Juenger

## Abstract

The leaf economics spectrum (LES) is hypothesized to result from a trade-off between resource acquisition and conservation. Yet few studies have examined the evolutionary mechanisms behind the LES, perhaps because most species exhibit relatively specialized leaf economics strategies. In a genetic mapping population of the phenotypically diverse grass *Panicum virgatum*, we evaluate two interacting mechanisms that may drive LES evolution: 1) genetic architecture, where multiple traits are coded by the same gene (pleiotropy) or by genes in close physical proximity (linkage), and 2) correlational selection, where selection acts non-additively on combinations of multiple traits. We found evidence suggesting that shared genetic architecture (pleiotropy) controls covariation between two pairs of leaf economics traits. Additionally, at five common gardens spanning 17 degrees of latitude, correlational selection favored particular combinations of leaf economics traits. Together, these results demonstrate how the LES can evolve within species.

## Introduction

Globally, plants exhibit a correlated suite of functional traits known as the worldwide leaf economics spectrum (Wright et al. 2004, Shipley et al. 2006, Reich 2014). This spectrum is hypothesized to result from a trade-off between resource acquisition in high resource environments and resource conservation in low resource environments (Donovan et al. 2011). The acquisitive end of this spectrum is characterized by a quick-return on investment strategy of short-lived leaves with high nutrient content and metabolic requirements. The conservative end is characterized by a slow-return on investment strategy of long-lived leaves with low nutrient content and metabolic requirements. While evidence for the worldwide leaf economics spectrum (LES) is highly consistent at the global scale (Wright et al. 2004, Díaz et al. 2016), results have been mixed at smaller spatial and taxonomic scales (e.g., Heberling and Fridley 2012, Edwards et al. 2014, Messier et al. 2017, Anderegg et al. 2018). These results have prompted several studies to examine how the LES originates, using either statistical (e.g., Shipley et al. 2006, Mason et al. 2016) or anatomical (e.g., Blonder et al. 2011, John et al. 2017, Onoda et al. 2017) perspectives. An important step toward resolving discrepancies among studies is to shift from describing the LES at different scales to examining the evolutionary forces that form the LES (Donovan et al. 2011). Two evolutionary mechanisms that can interact to promote or constraint LES evolution are correlational selection and genetic architecture.

Selection for favored trait combinations, or against maladapative trait combinations (i.e., correlational selection), is likely an important driver shaping functional trait correlations (Donovan et al. 2011). But the evolutionary responses to selection are also shaped by underlying genetic variation and covariance among traits (i.e., genetic architecture). Patterns of genetic covariance determine the possible trajectories of evolution and can constrain or facilitate the evolution of trait combinations (Walsh and Blows 2009). Thus, studying how adaptive syndromes like the LES evolve, requires accounting for both genetic architecture and correlational selection. Genetic architecture can arise from pleiotropy, where the same gene codes for multiple traits; from linkage, where genes coding for different traits are in close physical proximity and thus, are usually inherited together; or through linkage disequilibrium (LD), where alleles at different loci, and impacting different traits, are in statistical association (Lynch and Walsh 1998, Mackay 2001). Correlational selection occurs where selection acts on the covariance between two or more traits and those trait combinations reduce or enhance fitness (Lande and Arnold 1983). By favoring traits in combination, rather than isolation, correlational selection can promote phenotypic integration (Svensson et al. 2021), potentially explaining the presence of correlated leaf economics strategies. Moreover, the two mechanisms, correlational selection and genetic architecture (pleiotropy, linkage, LD), can interact to promote or constraint LES evolution (Sinervo and Svensson 2002). When selection favors certain trait combinations, but disadvantageous combinations are genetically linked, evolution of the LES will be constrained (Donovan et al. 2011). Over time though, selection can break down linkage and LD between these disadvantageous combinations (Guilherme Pereira and Des Marais 2020). Correlational selection can also promote the evolution of genetic linkage between traits (Svensson et al. 2021), generating and maintaining the genetic foundation for the leaf economics spectrum. When selection and genetic architecture simultaneously favor the same trait combinations, LES evolution can proceed more rapidly (Donovan et al. 2011).

For the LES to evolve, leaf economics traits must exhibit genetic variation. But leaf economics traits and correlations between traits can also be driven by environmental factors, like fertilization and drought (Sherrard and Maherali 2006, Fajardo and Siefert 2018, Ji et al. 2020). Conventional approaches to studying the LES cannot separate these genetic and environmental components (Swenson et al. 2020, Ahrens et al. 2021). However, the potential for LES evolution can be assessed using quantitative genetics techniques (Donovan et al. 2011). Metrics like heritability and genetic correlations, which quantify the genetic component of individual traits and the shared genetic contribution to correlations between traits, can show whether leaf economics traits possess sufficient genetic variation to evolve. Going further, genetic constraints on the LES can be identified through genetic mapping and selection on the LES can be tested using a selection gradient analysis (Donovan et al. 2011). A well-established genetic mapping technique—quantitative trait locus (QTL) mapping—involves crossing individuals from opposite ends of a phenotypic gradient, then testing for associations between the genome and phenotype (Mauricio 2001, Anderson and Mitchell-Olds 2011). Selection gradients can be estimated as the effect of a trait or trait combination on plant fitness and can be used to detect selection (Brodie et al. 1995, Conner and Hartl 2004, Caruso et al. 2020)

Recent efforts to quantify intraspecific variation in leaf economics traits has improved our understanding of the LES (Siefert et al. 2015, Messier et al. 2017, Anderegg et al. 2018). But, although intraspecific variation is required for evolution, these intraspecific studies rarely address how the LES evolves. To examine LES evolution, we leverage the strong ecotypic divergence in leaf economics strategies in *Panicum virgatum* L, a widespread C_4_ grass (Aspinwall et al. 2013). First, we examined the genetics of three leaf economics traits—leaf mass per area (LMA), leaf nitrogen content (N_MASS_), and photosynthetic rate (A_MASS_)—in a genetic mapping population of *P. virgatum* in a single common garden in central Texas. Then, to test for correlational selection on leaf economics traits, we examined the fitness of clones of these plants at five sites spanning 17 degrees of latitude in the central United States. Together, this allowed us to answer four questions: 1) are leaf economics traits under detectable genetic control? 2) do leaf economics traits covary? 3) is covariation amongst traits genetically driven? 4) do particular combinations of leaf economics traits increase fitness in the field?

## Methods

### Study system

*Panicum virgatum* is an ecologically and economically important species possessing large phenotypic variation in leaf economics traits. *Panicum virgatum* is common throughout central North American grasslands, occupying habitats that vary considerably in season length and mean annual temperature and precipitation (Lowry et al. 2014). To occupy these diverse habitats, *P. virgatum* has diverged into three ecotypes—upland, coastal, and lowland—with the coastal ecotype exhibiting characteristics intermediate between the other two (Casler 2012, Lovell et al. 2021). The upland ecotype typically occurs in northern regions with short growing seasons, and possesses an acquisitive strategy. The lowland ecotype, occurs in southern regions with long growing seasons, and possesses a conservative strategy (Aspinwall et al. 2013, Heckman et al. 2020).

### Experimental setup

This experimental design is described in detail by Milano et al. (2016) and (Lowry et al. 2019). In summary, a genetic mapping population was developed by crossing the southern lowland genotype AP13 × the northern upland genotype DAC and the southern lowland genotype WBC × the northern upland genotype VS16. A single F_1_ offspring from each of these two crosses was then crossed to produce a four-way outbred mapping population, which contained 400 full sibling F_2_ offspring. By recombining upland and lowland alleles, this cross generated F_2_ individuals that carry either two lowland alleles, two upland alleles, or one lowland and one upland allele at each locus. The F_2_ offspring were then clonally propagated in 3.8L pots at Brackenridge Field Laboratory, Austin, TX.

In February 2014, individuals of each genotype were transplanted into a common garden at Brackenridge Field Lab, where the soil is Yazoo sandy loam (Milano et al. 2016). The field was first covered with weed cloth and each individual was planted into a hole in the weed cloth. Plants were randomly assigned to locations in the field in a honeycomb design, with each plant located 1.25 m from its four nearest neighbors. To prevent edge effects, the field was surrounded with a border row of plants of the lowland genotype AP13.

### Leaf economics measurements

On 1-7 July 2014, we measured three leaf economics traits: mass-based photosynthetic capacity (A_MASS_), mass-based leaf nitrogen content (N_MASS_), and leaf mass per area (LMA). To calculate LMA, we measured the area of one penultimate fully expanded sun leaf per plant using a LI-3000C leaf area meter (LI-COR Biosciences, Lincoln, NE, USA), then dried leaves at 65°C to constant mass and weighed them. LMA is the ratio of leaf dry mass to leaf area. To calculate N_MASS_, these dried leaves were ground to a fine powder, then combusted in an elemental analyzer (Flash 2000 Organic Elemental NC Analyzer). To measure photosynthetic capacity, we enclosed two penultimate fully expanded sun leaves in the 2×3 cm cuvette of a LI-6400XT (LI-COR Biosciences, Lincoln, NE, USA) between 10:30 and 14:00. PAR was maintained within the cuvette at 1750 µmol m^-2^ s^-1^ using an actinic light source; chamber CO_2_ supply was set to 400 ppm, and leaf temperature and water vapor were allowed to track ambient conditions. We converted from an area-(A_AREA_) to mass-basis (A_MASS_) by dividing A_AREA_ by LMA.

### Fitness measurements

Clonally propagated individuals of each F_2_ line were planted in spring 2015 at five locations throughout central North America (Kingsville, TX – Brookings, SD) that spanned 17 degrees of latitude (details in Lowry et al 2019). Plantings at these five sites were identical in layout to the Austin, TX planting described above (see Lowry et al. 2019 for details). In 2017 and 2018, plants were harvested each fall ∼ 15 cm above ground level and aboveground biomass was dried and weighed. Biomasses measured in 2017 also appear in Lowry et al. (2019). Previous work has found that aboveground biomass is highly correlated with seed production in *P. virgatum* (Palik et al. 2016), making it a useful proxy for fitness (Lowry et al. 2019).

### Quantitative genetic and other statistical analyses

To assess whether leaf traits in *P. virgatum* represent a leaf economics spectrum, we performed standardized major axis (SMA) regression on each pair of leaf traits using the ‘sma’ function in smatr (Warton et al. 2012). SMA regression is commonly used in functional ecology because, unlike least squares regression, it assumes that both variables are measured with error (Warton et al. 2006).

To quantify the degree of genetic control of leaf economics traits, we calculated broad-sense heritability (H^2^) of leaf economics traits using the ‘mmer’ function in the sommer package (Covarrubias-Pazaran 2016). Heritability was the ratio of variance among lines (genetic variance, V_G_) to total variance (V_G_ + environmental variance, V_E_) in each trait (V_G_ / V_G_ + V_e_), calculated from a model that accounted for the relatedness among individuals by incorporating an additive genetic relatedness matrix. Throughout, we refer to V_G_ and H^2^ rather than additive genetic variance (V_A_) and narrow-sense heritability (h^2^) because additive and dominance genetic variance are confounded in this full-sib cross (Hill 2013).

To estimate the proportion of variance shared among leaf traits due to genetic causes (genetic correlations, r_g_), we performed a multivariate analysis using ‘mmer’. The model included all three leaf economics traits as responses with an additive genetic relatedness matrix as a random effect. To estimate the significance of these genetic correlations, we compared a model in which the genetic covariance between a pair of traits was estimated freely to a model in which the genetic covariance between those traits was constrained to 0 (i.e., no genetic covariance between traits was allowed) using a likelihood ratio test. A significant likelihood ratio test indicates that genetic covariance differs from 0.

### QTL mapping

We conducted QTL mapping on leaf economics traits using the qtl2 package (Broman et al. 2019). Methods for constructing the linkage map used in this analysis are described by Lovell et al. (2020). First, we assessed the likelihood that each genetic marker is associated with a phenotype of interest by using the ‘scan1’ function with a leave-one-chromosome-out kinship matrix. This provided log-odds (LOD) profiles for every marker in the genome. From these LOD profiles, we identified putative QTL using the ‘find_peak’ function with drop = 1.5 and peakdrop = 2.5 at a significance threshold of α = 0.15. We determined the LOD score corresponding to this significance threshold via permutation test with 1000 iterations using the ‘scan1perm’ function. We chose a relaxed significance threshold because a stricter threshold could hinder a primary goal of this study—to detect and evaluate QTL overlap among leaf traits. To maintain statistical rigor when evaluating putative QTL, we further assessed the significance and explanatory power of all QTL by fitting a multiple-QTL model for each phenotype using the ‘makeqtl’ and ‘fitqtl’ functions in the qtl package (Broman et al. 2003).

When QTL confidence intervals for two traits overlapped, we tested whether this overlap was due to pleiotropy (one QTL) or separate QTL using a pipeline from the qtl2pleio package (Boehm et al. 2019). To do this, we first used the ‘scan_pvl’ function to identify the region of QTL overlap, then used the ‘find_pleio_peak_tib’ function to identify the marker corresponding to the peak of the pleiotropy trace, then used the ‘boot_pvl’ function with 1000 iterations to perform a bootstrapped multivariate QTL scan that evaluates the evidence for separate QTL. In this analysis, the null hypothesis is one QTL (i.e., pleiotropy). Thus, P > 0.05 indicates a lack of evidence for separate QTL and is consistent with pleiotropy.

### Selection gradient analysis

To assess linear and non-linear selection on leaf economics traits in the field, we performed a selection gradient analysis. Standard selection gradient analysis pairs a fitness component (or fitness proxy) with traits measured on the same individuals (Lande and Arnold 1983). Here, however, we take a somewhat different approach: we measured leaf economics traits on plants at a single site, then pair these predictors with a fitness proxy measured on clonally propagated individuals at multiple sites. Because individuals grown at each site are genetic clones of the individuals measured for leaf economics traits, this is essentially a repeated-measures design. Importantly, we assume that plasticity and genotype-by-environment effects on LES traits are low. For selection gradients, we first mean-standardized genotype-level aboveground biomass separately for each of our five sites and two years by dividing each genotype’s biomass by the site-level mean biomass that year (Franklin and Morrissey 2017).

This results in a relativized proxy for fitness, such that average fitness at each site is equal to 1; above average fitness > 1; and below average fitness < 1. Plants that died during the study were retained for analysis with biomass = 0. At each of the three northern sites (Brookings, KBS, Columbia), only 0-4 plants were recorded as dead in any year. This corresponds with a mortality rate of 0-1.3%. At the two southern sites, mortality was higher: 5.7% and 16.2% in Kingsville in 2017 and 2018, respectively, and 0.3% and 8.2% in Austin in 2017 and 2018, respectively. We then examined how three mean-centered and variance-standardized (mean = 0, sd = 1) leaf economics traits impacted relative biomass in a single linear mixed model using the ‘lme’ function in nlme (Pinheiro et al. 2016). This model included main effects (i.e., linear directional selection), quadratic effects (i.e., non-linear stabilizing or disruptive selection), and all two-way interactions between these three traits (i.e., non-linear correlational selection) and interactions between all leaf economics predictors and site identity. Year of biomass measurement did not significantly interact with any other model parameters and was removed from the final model. To account for heteroscedasticity, this model included an identity variance structure (varIdent), which allows residual variance to differ by site.

To complement phenotypic selection gradients and circumvent the issue of measuring LES traits and biomass on different clonal propagates, we also performed genetic selection gradient analysis (Rausher 1992). To do this, we first calculated genomic best linear unbiased predictors (GBLUPs) for LES traits and biomass using the ‘mmer’ model described above for estimating heritability. GBLUPs represent the additive genetic contribution to a given phenotype. We then performed selection gradient analysis with GBLUPs instead of phenotypic values using the ‘gls’ function. This approach assesses only the genetic relationship between traits, which more accurately describes the evolutionary pressure on LES traits.

## Results

### Are leaf economics traits under detectable genetic control?

All three traits exhibited moderate broad-sense heritability (H^2^; Table 1). Of the three, A_MASS_ had the lowest heritability (H^2^ = 0.26), while LMA and N_MASS_ had similar, and higher, heritability (H^2^ = 0.51 and 0.46, respectively). For each trait, we detected 4-5 significant QTL (Table 2), which combined to explain 19-33% of variation in leaf economics traits (Table 1). Together, these results confirm that the leaf economics traits evaluated here have a substantial genetic basis.

**Table 1.** Genetic basis of three leaf economics traits. H^2^ is broad-sense heritability [V_G_ / (V_G_ + V_e_)] ± standard error; variance explained by QTL is calculated from a multiple-QTL model using all identified putative QTL as predictors of each trait; proportion of heritable variation explained by QTL is the ratio of variance explained by QTL to H^2^. LMA is leaf mass per area (g m^-2^); N_MASS_ is mass-based leaf nitrogen content (%); A_MASS_ is mass-based leaf photosynthetic rate (µmol C g^-1^ s^-1^).

| | $H^2$ | Variance explained<br>by QTL | Proportion of heritable<br>variation explained by QTL |
| --- | --- | --- | --- |
| LMA | $0.51 \pm 0.07$ | 0.33 | 0.65 |
| $N_{\text{MASS}}$ | $0.46 \pm 0.08$ | 0.24 | 0.52 |
| $A_{\text{MASS}}$ | $0.26 \pm 0.08$ | 0.19 | 0.73 |

**Table 2.** Significant QTL markers for each leaf economics trait detected at a LOD threshold of α = 0.15. P values are based on F tests calculated by dropping one QTL at a time from a multiple-QTL model that included all putative QTL for each trait.

| Trait | Marker | LOD | P |
| --- | --- | --- | --- |
| LMA | Chr2N@56.080999 | 3.972 | < 0.001 |
|  | Chr5N@4.933063 | 7.814 | < 0.001 |
|  | Chr8N@13.652308 | 3.873 | < 0.001 |
|  | Chr9K@12.433663 | 10.359 | < 0.001 |
|  | Chr9N@79.37425 | 4.173 | < 0.001 |
| A<sub>MASS</sub> | Chr2K@11.849289 | 6.079 | < 0.001 |
|  | Chr2K@59.819311 | 4.58 | 0.017 |
|  | Chr8K@11.338989 | 4.771 | < 0.001 |
|  | Chr9K@13.273228 | 3.896 | 0.002 |
| N<sub>MASS</sub> | Chr2K@10.283923 | 4.164 | < 0.001 |
|  | Chr2K@61.75027 | 5.601 | < 0.001 |
|  | Chr3N@16.536961 | 8.592 | < 0.001 |
|  | Chr5K@21.5219 | 3.929 | < 0.001 |

### Do leaf economics traits covary phenotypically?

In F_2_ individuals, there were significant bivariate relationships amongst all leaf economics traits. LMA covaried negatively with N_MASS_ (R^2^ = 0.06, P < 0.001; Fig. 1a) and with A_MASS_ (R^2^ = 0.29, P < 0.001; Fig. 1b), which is consistent with global LES relationships. Similarly, N_MASS_ and A_MASS_ covaried positively (R^2^ = 0.06, P < 0.001; Fig. 1c). Thus, in this system, leaf economics traits significantly, although sometimes weakly, covary phenotypically.

**Fig 1.**
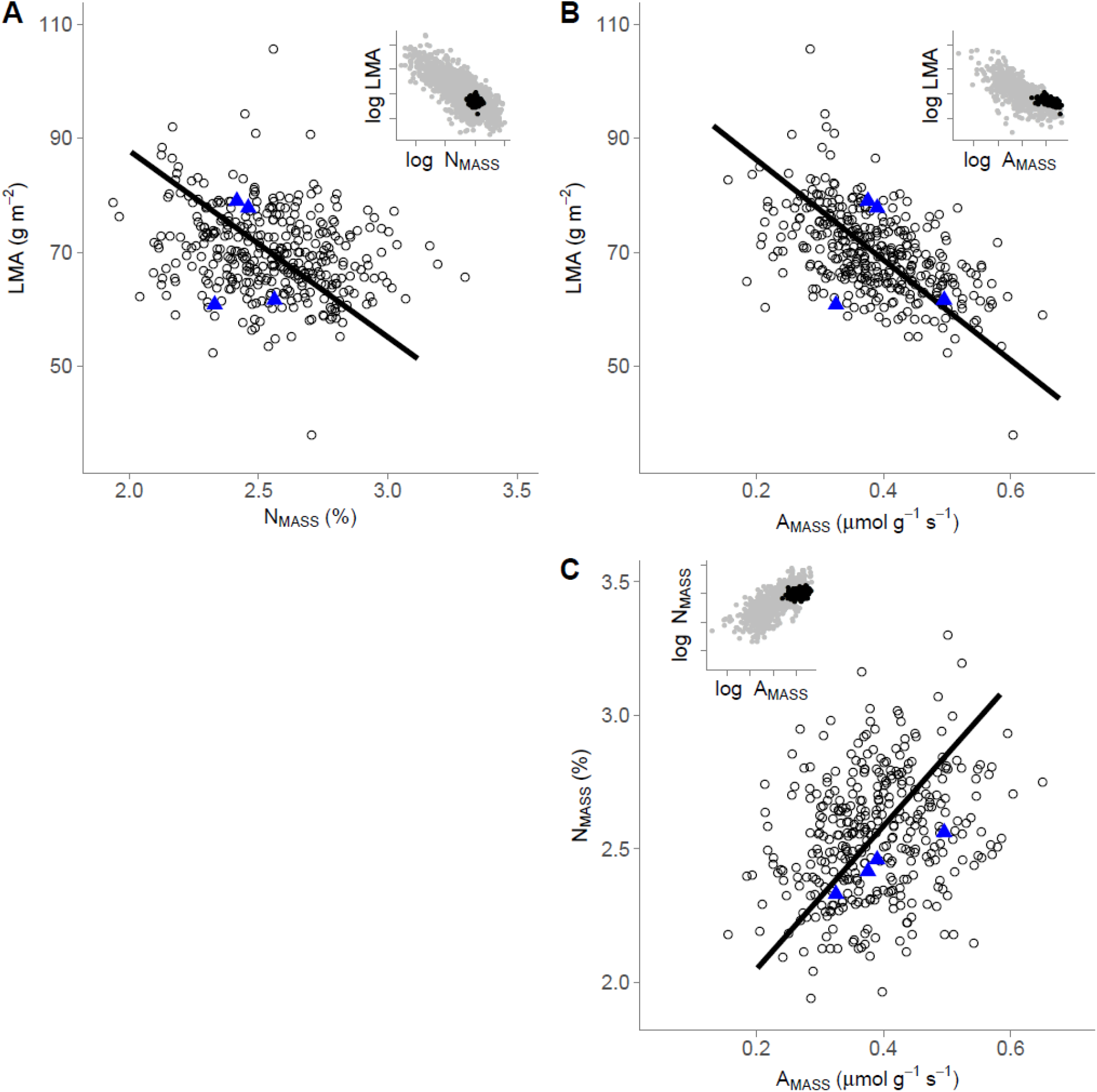
Relationship between **A)** leaf mass-based nitrogen (N_MASS_; %) and leaf mass per area (LMA; g m^-2^); **B)** leaf mass-based photosynthetic rate (A_MASS_; µmol C g^-1^ s^-1^) and LMA; **C)** A_MASS_ and N_MASS_ calculated using standardized major axis regression. Black dots are *P. virgatum* F_2_ individuals; blue triangles are F_0_ line means (not included in SMA line fit). Inset plots show the location of *P. virgatum* F_2_ individuals (black dots) relative to the worldwide leaf economics spectrum (grey dots; from Wright et al. 2004) on a log_10_-log_10_ scale.

### Is covariation amongst traits genetically driven?

We detected 13 QTL for leaf economics traits, all of which were highly significant predictors of LES traits in multiple-QTL models (P < 0.05; Table 2). These 13 QTL included three pairs of colocalized QTL (i.e., confidence intervals for the QTL overlapped; Fig. 2a): LMA-A_MASS_ on Chr9K and A_MASS_-N_MASS_ at two locations on Chr2K. For all three pairs of colocalized QTL, our results are consistent with pleiotropy (i.e., one QTL for both traits) because we did not find evidence that these colocalized QTL were produced by distinct loci (P = 0.14, P = 0.254, and P = 0.429, respectively; Fig. 2b). Further supporting pleiotropy, marker regression showed that genotype effects at colocalized QTL were in the same direction for both traits (i.e., both conservative or both acquisitive). For instance, individuals possessing two upland alleles on Chr2K@12 had lower A_MASS_ and N_MASS_ than other genotypes. Similarly, at the QTL on Chr9K, individuals with two upland alleles had significantly higher A_MASS_ and significantly lower LMA than other genotypes. In total, these results show that leaf economics traits exhibit some degree of genetic coordination.

**Fig 2.**
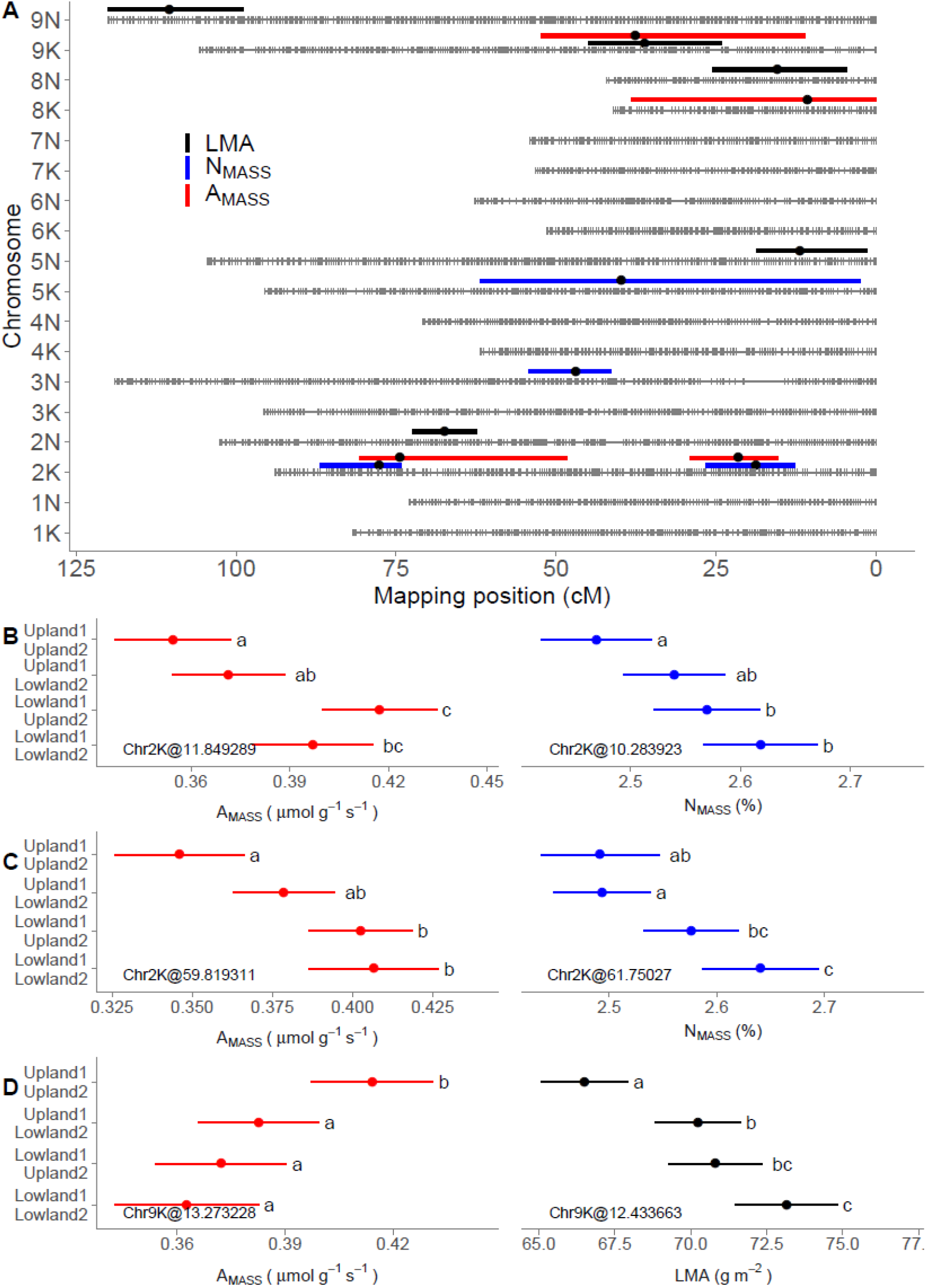
**A** Location of QTL for three leaf economics traits, LMA, N_MASS_, and A_MASS_. Point estimates are the location of highest LOD score and confidence intervals are the region within a 1.5 LOD drop. **B** Effects of genotype on leaf economics traits at three colocalized QTL markers (i.e., markers with overlapping confidence intervals) calculated using ordinary least squares regression. Genotypes at each QTL are a consequence of recombination in the experimental cross, resulting in four possible combinations of alleles. In each panel, lowland1 and lowland2 denote alleles inherited from genotypes AP13 and WBC, respectively; upland1 and upland2 denote alleles inherited from genotypes DAC and VS16, respectively. Error bars represent 95% confidence intervals; shared letters indicate no significant difference between genotypes. For all colocialized QTL pairs, genotype effects are in a consistent direction (i.e., each genotype exhibits conservative or acquisitive values of both traits) and are consistent with pleiotropy. See Fig. 1 legend for units and abbreviations.

Despite seeing colocalized QTL for LMA-A_MASS_ and N_MASS_-A_MASS_, a significant genetic correlation existed only between LMA and A_MASS_ (r_g_ = -0.72, χ^2^ = 15.87, P < 0.001; Table 3). Neither LMA-N_MASS_ (r_g_ = -0.12, χ^2^ = 0.79, P = 0.37) nor A_MASS_-N_MASS_ (r_g_ = 0.26, χ^2^ = 1.11, P = 0.29) had significant genetic correlations. These contrasting effects may be related to the directionality of QTL for the three LES traits. All five significant QTL for LMA were in the same direction and consistent with a conservative leaf economics strategy among the lowland ecotype and an acquisitive strategy among the upland ecotype (i.e., individuals with both lowland alleles had high LMA and individuals with both upland alleles had low LMA; Figs. 2d, S2a-d). Conversely, three of the four N_MASS_ QTL and two of the four A_MASS_ QTL behaved opposite to LES predictions: individuals with both upland alleles had low N_MASS_ and low A_MASS_, while individuals with both lowland alleles had high N_MASS_ and high A_MASS_ (Figs 2b-c, S2g). When QTL differ in directionality, as LMA and N_MASS_ did, it may weaken the aggregate genetic correlations (Gromko 1995). Similarly, despite detecting two colocalized QTL for A_MASS_ and N_MASS_, these pairs of colocalized QTL were all in the opposite direction of typical leaf economics strategies, which could also weaken genetic correlations.

**Table 3.** Genetic correlations between leaf economics traits. LMA is leaf mass per area (g m^-2^); N_MASS_ is mass-based leaf nitrogen content (%); A_MASS_ is mass-based leaf photosynthetic rate (µmol C g^-1^ s^-1^).

| | LMA | $A_{\text{MASS}}$ | $N_{\text{MASS}}$ |
| --- | --- | --- | --- |
| LMA | --- |  |  |
| $A_{\text{MASS}}$ | **-0.76** (0.12) | --- | |
| $N_{\text{MASS}}$ | -0.16 (0.17) | 0.26 (0.21) | --- |

### Do particular combinations of leaf economics traits increase fitness in the field?

There was a significant interactive effect of LMA and N_MASS_ on relative biomass across all sites (γ_*ij*_ = -0.05; P = 0.015; Table S2, S3; Fig. 3a). The relative biomass of individuals possessing conservative values of both traits (i.e., high LMA, low N_MASS_) was higher than that of individuals with acquisitive values of both traits (i.e., low LMA, high N_MASS_). Both conservative and acquisitive individuals had greater relative biomass than individuals with mismatched trait combinations (high LMA, high N_MASS_ or low LMA, low N_MASS_). This correlational selection for particular combinations of LMA and N_MASS_ could promote LES evolution. Additionally, when controlling for other trait values, LMA had a marginally significant direct positive effect on relative biomass across all sites (LMA: β = 0.03; P = 0.081; Table S2, S3).

**Fig 3.**
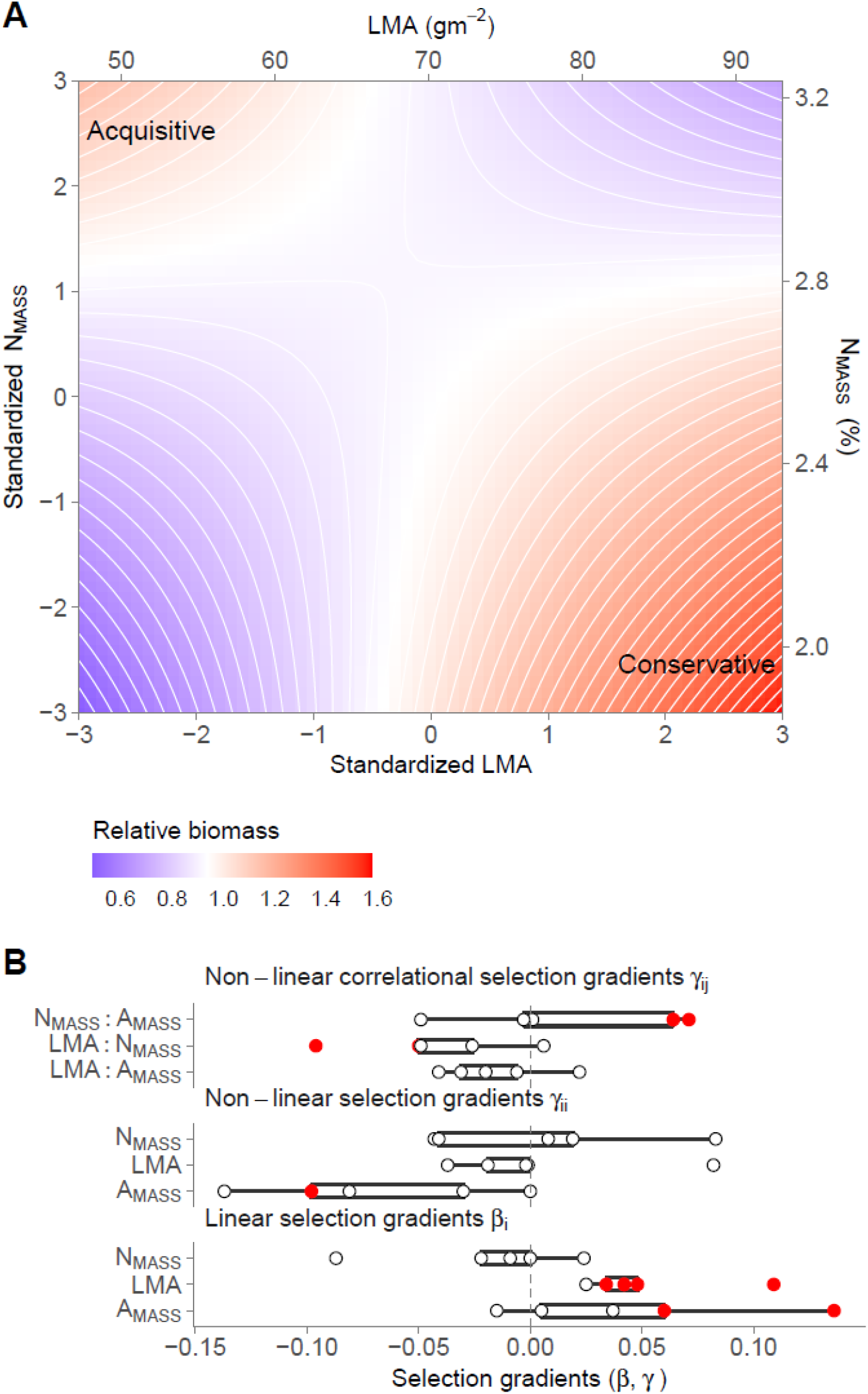
**A** Effects of standardized LMA and N_MASS_ (transformed such that across all plants, mean = 0 and sd = 1) on relative biomass while controlling for A_MASS_. Following (Stinchcombe et al. 2008), model-derived quadratic parameter estimates are doubled. **B** Selection gradients estimated at each site. Linear selection gradients (β_i_) are main effects; non-linear selection gradients (γ_ii_) are quadratic effects (model-derived estimates were doubled); non-linear correlational selection gradients (γ_ij_) are interactive effects between two leaf economics traits. Red points denote significant effect. See Fig. 1 legend for units and abbreviations.

Selection gradients differed among sites, but not systematically by latitude (Table S3, S4; Fig. 3b). Directional selection favored high LMA plants at four of five sites (Site × LMA: P = 0.42; KBSM: β = 0.06, P = 0.014; CLMB: β = 0.04, P = 0.095; PKLE: β = 0.09, P = 0.046; KING: β = 0.12, P = 0.063; Table S3, S4), and high A_MASS_ plants at two of five sites (Site × A_MASS_: P = 0.008; KBSM: β = 0.06, P = 0.015; KING: β = 0.13, P = 0.053; Table S3, S4). Selection favoring higher values of both LMA and A_MASS_ combined with a negative genetic correlation between the two traits would generate a genetic constraint on the evolution of the LES in *P. virgatum*. Finally, correlational selection favored matched phenotypic combinations of LMA and N_MASS_ at two sites (Site × LMA × N_MASS_: P = 0.27; BRKG: P = 0.059; PKLE: P = 0.045; Table S4; Fig. S2) and matched combinations of A_MASS_ and N_MASS_ at two other sites (Site × N_MASS_ × A_MASS_: P = 0.009; KBSM: P = 0.007; CLMB: P = 0.018; Table S4; Fig. S3).

Genetic selection gradients largely supported these results: LMA × N_MASS_ and N_MASS_ × A_MASS_ interacted to impact biomass among GBLUPs (P = 0.080 and P = 0.032, respectively; Table S5; Fig 4). Together, these results demonstrate that correlational selection on particular leaf economics trait combinations can occur in the field and is genetically based.

**Fig 4.**
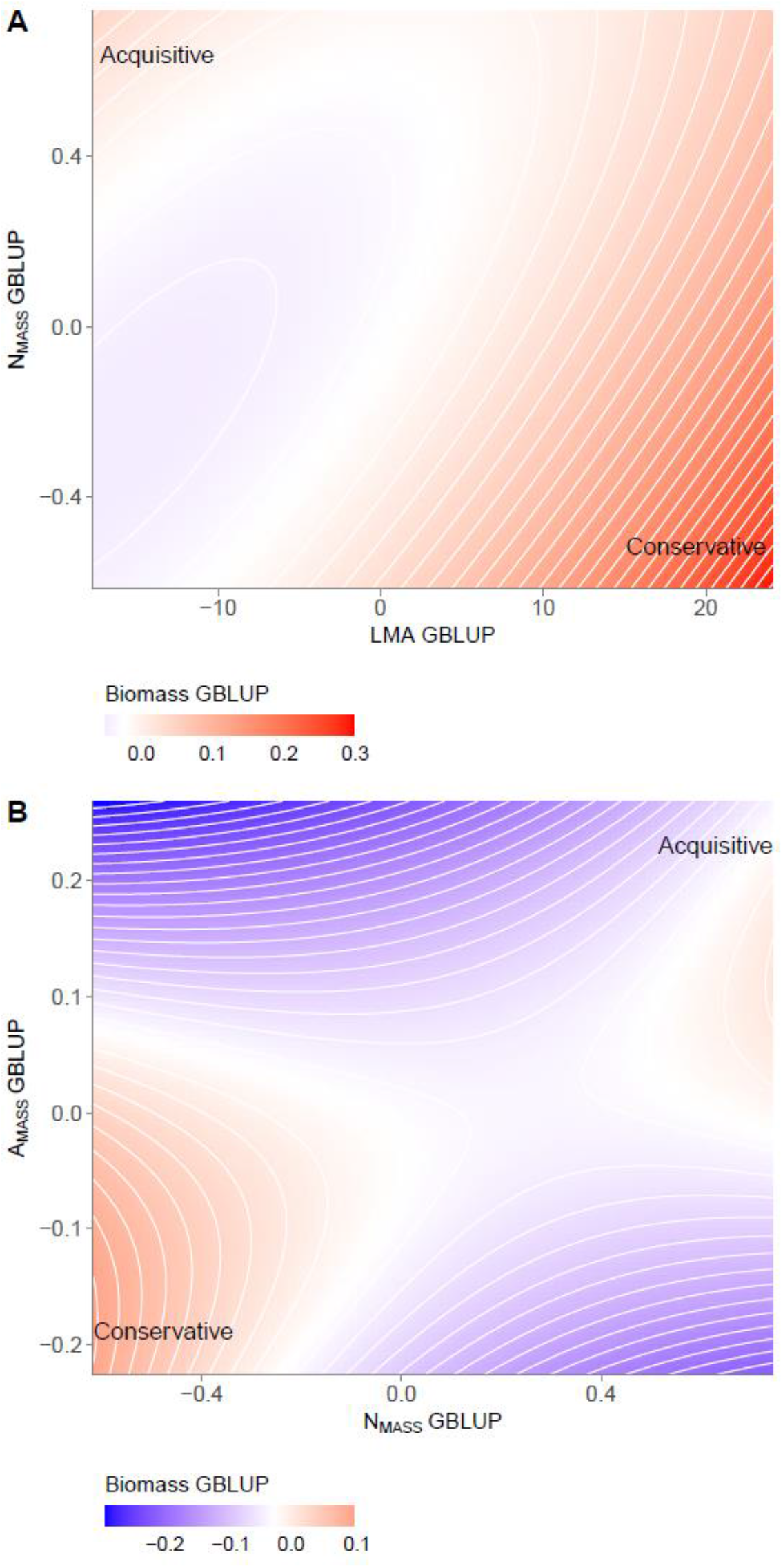
**A** Effects of LMA and N_MASS_ genomic linear unbiased predictors (GBLUPs) on plant biomass GBLUPs **B** Effects of N_MASS_ and A_MASS_ GBLUPs on plant biomass GBLUPs. GBLUPs were calculated separately for each trait from a mixed model that accounted for the relatedness among individuals by incorporating an additive genetic relatedness matrix

## Discussion

We found evidence that the LES exists in *P. virgatum*, and may be driven by a combination of genetic architecture and correlational selection. These results provide the strongest evidence to date for the evolution of the LES within a species. Overall, we found that correlational selection and genetic architecture generally promote the hypothesized leaf economics strategies, but that some genetic constraints may limit LES evolution in this system.

Few studies have explored the mechanisms driving the evolution of leaf economics strategies (Donovan et al. 2011). This results, in part, from the fact that most species, and even many genera, occupy a small portion of the LES (Edwards et al. 2014). Because most species possess relatively specialized leaf economics strategies, they typically lack sufficient variation for genetic constraints and correlational selection to promote LES evolution (Agrawal 2020).

But, when species exhibit distinct ecotypes or occupy a broad range of habitats, it is possible to evolve a clear LES at lower taxonomic levels, as we observe in *P. virgatum*. Thus, species like *P. virgatum*, that do possess distinct leaf economic strategies, offer a unique opportunity to evaluate microevolution of the LES. By crossing genotypes that exhibit divergent leaf economics strategies, we can break up these leaf economics strategies and examine how selection acts on combinations of traits that rarely occur naturally. This is especially valuable because it is impossible to independently manipulate two or more LES traits in a macroevolutionary context (Shipley et al. 2006).

Our approach is not the only valuable way to study the evolution of the LES. One important method for understanding genetic correlations between traits is artificial selection on extreme trait combinations. Donovan et al. (2011) advocate for selection on high N_MASS_ – low A_MASS_ lines (and vice versa) for multiple generations to determine whether a positive correlation between N_MASS_ and A_MASS_ can be broken up. Another way to evaluate selection on leaf economics strategies is to grow plants under different resource conditions, then examine how fitness changes. For instance, to test the hypothesis that acquisitive strategies are favored under high resource conditions and conservative strategies are favored under low resource conditions, one could evaluate the fitness of conservative and acquisitive genotypes along an experimental nutrient gradient. This approach could show not only whether the LES is present within a species, but also, whether it should evolve through the hypothesized acquisition-conservation trade-off.

The selective pressures acting upon individual species may differ from those that produce the LES among species, potentially limiting the evolution of the LES within species (Agrawal 2020). For instance, when species specialize in particular habitats or evolve distinct ecotypes adapted to habitats differing in water availability, these selective pressures may not align with the LES. Thus, LES evolution within species may be rare (Agrawal 2020). In *P. virgatum*, the evolution of distinct leaf economics strategies may be driven by large differences in growing season lengths across its range. The northern upland ecotype, which exhibits an acquisitive strategy (e.g., smaller, narrower leaves with higher N_MASS_ and A_MASS_ and lower LMA), may have evolved to take advantage of the relatively short growing season in the northern US and Canada. Short growing seasons may promote a quick return-on-investment strategy of low tissue construction costs and high photosynthetic capacity (Baird et al. 2021). In the southern habitat typical of the lowland ecotype, the long growing season may select for longer-lived leaves with higher LMA.

Season length as a driver of leaf economics strategies may be particularly important if divergent selection pressures in the northern and southern portions of *P. virgatum*’s range drive selection for different leaf economics strategies. All five putative QTL for LMA exhibited divergence in the same direction—plants with lowland alleles exhibited higher LMA than plants with upland alleles—suggesting that directional selection may have promoted divergence in LMA across the range of *P. virgatum* (Milano et al. 2016). Although all significant QTL for LMA had the same directionality, QTL for A_MASS_ and N_MASS_ did not. This suggests that selection on these two traits may depend on infrequent, severe abiotic or biotic stress, such as cold, drought, or high consumer pressure (e.g., some strategies may be advantageous only under severe stress), or stabilizing selection that maintains moderate values of the traits. This may slow the evolution of leaf economics strategies in *P. virgatum*. Evolution of leaf economics strategies may also be constrained by the LMA-A_MASS_ relationship: high LMA and high A_MASS_ were both favored in selection gradient analysis, but the strong negative genetic correlation between these traits should limit the ability of this combination to evolve. On the other hand, constraints on the evolution of N_MASS_-LMA should be minimal. These traits exhibited a weak negative genetic correlation and correlational selection in the same direction (i.e., high LMA and low N_MASS_ are favored). Together, these results suggest that the conservative strategy should be favored under most circumstances, but the acquisitive strategy (low LMA, high N_MASS_) could be favored under more extreme conditions that did not occur over the duration of this study. While it is impractical to follow these individuals over their entire lifetimes, future studies that incorporate climatic extremes, either naturally or experimentally, could better determine the conditions that favor the acquisitive strategy in *P. virgatum*.

Our results showed some similarities with the small number of previous studies on this topic. Like Donovan et al. (2011), we found that genetic correlations between leaf economics traits were variable and while directional selection occurred on many leaf economics traits in the studies they surveyed, selection differentials and gradients were not consistent between traits or across studies. However, due to limitations in the data available to Donovan et al. (2011), there were also some notable differences between our studies. For instance, we found evidence for pleiotropic loci controlling both N_MASS_-A_MASS_ and A_MASS_-LMA and a potential genetic constraint on the correlated evolution of LMA and A_MASS_. Our results are also somewhat consistent with recent work on the LES in *Arabidopsis thaliana*, which showed that pleiotropic loci control some leaf economics correlations (Vasseur et al. 2012, Hanemian et al. 2020) and that leaf economics strategies differ across the geographic range of the species, possibly due to climate-driven selection (Sartori et al. 2019).

### Limitations

This study had several limitations. First, we did not measure leaf economics traits and fitness on the same plants. This would be important if plasticity in leaf economics traits is high, particularly if genotypes differ in plasticity (i.e., genotype-by-environment interactions). In a previous study using different genotypes of *P. virgatum*, we found significant genotype-by-environment effects on N_MASS_, but not on leaf dry matter content, which is highly correlated with LMA (Heckman et al. 2020). Given the low heritability of A_MASS_, this trait is likely to exhibit higher plasticity, although it is unclear whether A_MASS_ should exhibit strong GxE. Because genetic selection gradients were largely consistent with phenotypic selection gradients, plasticity or environmental correlations in traits should be a minor concern. However, genetic selection gradients could still be biased by large genotype-by-environment effects. Second, we use biomass as a proxy for fitness (Franklin and Morrissey 2017). While biomass production is highly correlated with seed set (Lowry et al. 2019), we cannot assess fitness over the entire lifespan of this relatively long-lived species. Third, because most ecologically important quantitative traits are polygenic (Barghi et al. 2020), we probably failed to detect many important QTL. This is made clear by the fact that significant QTL explained only ∼50% of the genetic variation in these traits. Thus, these QTL results should be considered a hypothesis-generating tool. Future work could examine in new populations whether the same QTL are detected. Moreover, future studies could further explore candidate genes in the overlapping QTL regions that suggest pleiotropy.

## Conclusions

The worldwide LES is one of the most striking patterns in plant ecology, yet little work has been done to examine how it evolves. Here, we show that the LES can evolve within a widespread grass species through a combination of genetically linked traits and correlational selection favoring individuals possessing particular LES combinations. While it is unclear whether evolution of the LES in *P. virgatum* is driven by the same resource conservation-acquisition trade-off that is hypothesized to underlie the worldwide LES, this system provides a rare opportunity to address this longstanding hypothesis in a tractable system.

## Acknowledgments

We thank Quinn Hiers and Albina Khasanova for help with data collection. Elizabeth Alger, Tina Arredondo, Nicole Carrabba, Elena Pinaroc, Maya Rao, Maria Villalpando, and Heather Yang also provided field assistance. Alice MacQueen and other members of the Juenger lab provided helpful comments on a previous draft of the manuscript. This work was supported by NSF PGRP IOS 0922457 and IOS 1444533. This research was supported by the Office of Science (BER), U.S. Department of Energy, Grant no. DE-SC0014156. USDA ARS is an Equal Opportunity Employer. Mention of trade names or commercial products in this publication does not imply recommendation or endorsement by the USDA.

## Author contributions

RWH analyzed the data and led writing with input from TEJ and PAF; TEJ conceived the experiment; JEB and BEC collected data and contributed to writing.

## References

Agrawal, A. A. 2020. A scale-dependent framework for trade-offs, syndromes, and specialization in organismal biology. Ecology 101:e02924.

Ahrens, C. W., P. D. Rymer, and D. T. Tissue. 2021. Intra-specific trait variation remains hidden in the environment. New Phytologist 229:1183–1185.

Anderegg, L. D. L., L. T. Berner, G. Badgley, M. L. Sethi, B. E. Law, J. HilleRisLambers, and J. Penuelas. 2018. Within-species patterns challenge our understanding of the leaf economics spectrum. Ecology Letters 21:734–744.

Anderson, J. T., and T. Mitchell-Olds. 2011. Ecological genetics and genomics of plant defences: evidence and approaches. Functional Ecology 25:312–324.

Aspinwall, M. J., D. B. Lowry, S. H. Taylor, T. E. Juenger, C. V. Hawkes, M.-V. V. Johnson, J. R. Kiniry, and P. A. Fay. 2013. Genotypic variation in traits linked to climate and aboveground productivity in a widespread C_4_grass: evidence for a functional trait syndrome. New Phytologist 199:966–980.

Baird, A. S., et al. 2021. Developmental and biophysical determinants of grass leaf size worldwide. Nature 592:242–247.

Barghi, N., J. Hermisson, and C. Schlötterer. 2020. Polygenic adaptation: a unifying framework to understand positive selection. Nature Reviews Genetics 21:769–781.

Blonder, B., C. Violle, L. P. Bentley, and B. J. Enquist. 2011. Venation networks and the origin of the leaf economics spectrum. Ecology Letters 14:91–100.

Boehm, F. J., E. J. Chesler, B. S. Yandell, and K. W. Broman. 2019. Testing pleiotropy vs. separate QTL in multiparental populations. G3 Genes|Genomes|Genetics 9:2317–2324.

Brodie, E. D., A. J. Moore, and F. J. Janzen. 1995. Visualizing and quantifying natural selection. Trends in Ecology & Evolution 10:313–318.

Broman, K. W., D. M. Gatti, P. Simecek, N. A. Furlotte, P. Prins, ś. Sen, B. S. Yandell, and G. A. Churchill. 2019. R/qtl2: Software for Mapping Quantitative Trait Loci with High-Dimensional Data and Multiparent Populations. Genetics 211:495–502.

Broman, K. W., H. Wu, ś. Sen, and G. A. Churchill. 2003. R/qtl: QTL mapping in experimental crosses. Bioinformatics 19:889–890.

Caruso, C. M., C. M. Mason, and J. S. Medeiros. 2020. The evolution of functional traits in plants: is the giant still sleeping? International Journal of Plant Sciences 181:1–8.

Casler, M. D. 2012. Switchgrass Breeding, Genetics, and Genomics. Pages 29–53 in A. Monti, editor. Switchgrass: A Valuable Biomass Crop for Energy. Springer London, London.

Conner, J. K., and D. L. Hartl. 2004. A primer of ecological genetics. Sinauer Associates Incorporated.

Covarrubias-Pazaran, G. 2016. Genome-assisted prediction of quantitative traits using the R package sommer. Plos One 11:e0156744.

Díaz, S., et al. 2016. The global spectrum of plant form and function. Nature 529:167–171.

Donovan, L. A., H. Maherali, C. M. Caruso, H. Huber, and H. de Kroon. 2011. The evolution of the worldwide leaf economics spectrum. Trends in Ecology & Evolution 26:88–95.

Edwards, E. J., D. S. Chatelet, L. Sack, and M. J. Donoghue. 2014. Leaf life span and the leaf economic spectrum in the context of whole plant architecture. Journal of Ecology 102:328–336.

Fajardo, A., and A. Siefert. 2018. Intraspecific trait variation and the leaf economics spectrum across resource gradients and levels of organization. Ecology 99:1024–1030.

Franklin, O. D., and M. B. Morrissey. 2017. Inference of selection gradients using performance measures as fitness proxies. Methods in Ecology and Evolution 8:663–677.

Gromko, M. H. 1995. Unpredictability of correlated response to selection: pleiotropy and sampling interact. Evolution 49:685–693.

Guilherme Pereira, C., and D. L. Des Marais. 2020. The genetic basis of plant functional traits and the evolution of plant-environment interactions. International Journal of Plant Sciences 181:56–74.

Hanemian, M., F. Vasseur, E. Marchadier, E. Gilbault, J. Bresson, I. Gy, C. Violle, and O. Loudet. 2020. Natural variation at FLM splicing has pleiotropic effects modulating ecological strategies in Arabidopsis thaliana. Nature Communications 11:4140.

Heberling, J. M., and J. D. Fridley. 2012. Biogeographic constraints on the world-wide leaf economics spectrum. Global Ecology and Biogeography 21:1137–1146.

Heckman, R. W., A. R. Khasanova, N. S. Johnson, S. Weber, J. E. Bonnette, M. J. Aspinwall, L. G. Reichmann, T. E. Juenger, P. A. Fay, and C. V. Hawkes. 2020. Plant biomass, not plant economics traits, determines responses of soil CO_2_efflux to precipitation in the C_4_grass Panicum virgatum. Journal of Ecology 108:2095–2106.

Hill, W. G. 2013. On estimation of genetic variance within families using genome-wide identity-by-descent sharing. Genetics Selection Evolution 45:32.

Ji, W., S. E. LaZerte, M. J. Waterway, and M. J. Lechowicz. 2020. Functional ecology of congeneric variation in the leaf economics spectrum. New Phytologist 225:196–208.

John, G. P., C. Scoffoni, T. N. Buckley, R. Villar, H. Poorter, and L. Sack. 2017. The anatomical and compositional basis of leaf mass per area. Ecology Letters 20:412–425.

Lande, R., and S. J. Arnold. 1983. The measurement of selection on correlated characters. Evolution:1210–1226.

Lovell, J., A. Healey, J. Schmutz, and T. Juenger. 2020. Switchgrass v5 4-way (AP13 x DAC, WBC x VS16) genetic map. Dryad.

Lovell, J. T., et al. 2021. Genomic mechanisms of climate adaptation in polyploid bioenergy switchgrass. Nature 590:438–444.

Lowry, D. B., K. D. Behrman, P. Grabowski, G. P. Morris, J. R. Kiniry, and T. E. Juenger. 2014. Adaptations between ecotypes and along environmental gradients in Panicum virgatum. American Naturalist 183:682–692.

Lowry, D. B., et al. 2019. QTL × environment interactions underlie adaptive divergence in switchgrass across a large latitudinal gradient. Proceedings of the National Academy of Sciences 116:12933.

Lynch, M., and B. Walsh. 1998. Genetics and analysis of quantitative traits.

Mackay, T. F. C. 2001. The genetic architecture of quantitative traits. Annual Review of Genetics 35:303–339.

Mason, C. M., E. W. Goolsby, D. P. Humphreys, and L. A. Donovan. 2016. Phylogenetic structural equation modelling reveals no need for an ‘origin’ of the leaf economics spectrum. Ecology Letters 19:54–61.

Mauricio, R. 2001. Mapping quantitative trait loci in plants: uses and caveats for evolutionary biology. Nature Reviews Genetics 2:370–381.

Messier, J., B. J. McGill, B. J. Enquist, and M. J. Lechowicz. 2017. Trait variation and integration across scales: is the leaf economic spectrum present at local scales? Ecography 40:685–697.

Milano, E. R., D. B. Lowry, and T. E. Juenger. 2016. The genetic basis of upland/lowland ecotype divergence in switchgrass (Panicum virgatum). G3 Genes|Genomes|Genetics 6:3561–3570.

Onoda, Y., I. J. Wright, J. R. Evans, K. Hikosaka, K. Kitajima, Ü. Niinemets, H. Poorter, T. Tosens, and M. Westoby. 2017. Physiological and structural tradeoffs underlying the leaf economics spectrum. New Phytologist 214:1447–1463.

Palik, D. J., A. A. Snow, A. L. Stottlemyer, M. N. Miriti, and E. A. Heaton. 2016. Relative performance of non-local cultivars and local, wild populations of Switchgrass (Panicum virgatum) in competition experiments. Plos One 11:e0154444.

Pinheiro, J., D. Bates, S. DebRoy, and D. Sarkar. 2016. nlme: linear and nonlinear mixed effects models. R package version 3. 1–127.

Rausher, M. D. 1992. The meausrement of selection on quantitative traits: biases due to environmental covariances between traits and fitness. Evolution 46:616–626.

Reich, P. B. 2014. The world-wide ‘fast–slow’ plant economics spectrum: a traits manifesto. Journal of Ecology 102:275–301.

Sartori, K., et al. 2019. Leaf economics and slow-fast adaptation across the geographic range of Arabidopsis thaliana. Scientific Reports 9:10758.

Sherrard, M. E., and H. Maherali. 2006. The adaptive significance of drought escape in Avena barbata, an annual grass. Evolution 60:2478.

Shipley, B., M. J. Lechowicz, I. Wright, and P. B. Reich. 2006. Fundamental trade-offs generating the worldwide leaf economics spectrum. Ecology 87:535–541.

Siefert, A., et al. 2015. A global meta-analysis of the relative extent of intraspecific trait variation in plant communities. Ecology Letters 18:1406–1419.

Sinervo, B., and E. Svensson. 2002. Correlational selection and the evolution of genomic architecture. Heredity 89:329–338.

Stinchcombe, J. R., A. F. Agrawal, P. A. Hohenlohe, S. J. Arnold, and M. W. Blows. 2008. Estimating nonlinear selection gradients using quadratic regression coefficients: double or nothing? Evolution 62:2435–2440.

Svensson, E. I., et al. 2021. Correlational selection in the age of genomics. Nature Ecology & Evolution 5:562–573.

Swenson, N. G., S. J. Worthy, D. Eubanks, Y. Iida, L. Monks, K. Petprakob, V. E. Rubio, K. Staiger, and J. Zambrano. 2020. A reframing of trait–demographic rate analyses for ecology and evolutionary biology. International Journal of Plant Sciences 181:33–43.

Vasseur, F., C. Violle, B. J. Enquist, C. Granier, and D. Vile. 2012. A common genetic basis to the origin of the leaf economics spectrum and metabolic scaling allometry. Ecology Letters 15:1149–1157.

Walsh, B., and M. W. Blows. 2009. Abundant genetic variation + strong selection = multivariate genetic constraints: a geometric view of adaptation. Annual Review of Ecology, Evolution, and Systematics 40:41–59.

Warton, D. I., R. A. Duursma, D. S. Falster, and S. Taskinen. 2012. smatr 3– an R package for estimation and inference about allometric lines. Methods in Ecology and Evolution 3:257–259.

Warton, D. I., I. J. Wright, D. S. Falster, and M. Westoby. 2006. Bivariate line-fitting methods for allometry. Biological Reviews 81:259–291.

Wright, I. J., et al. 2004. The worldwide leaf economics spectrum. Nature 428:821–827.

