## Supplementary Information for "Correlational selection and genetic architecture promote the leaf economics spectrum in a perennial grass"

The following Supporting Information is available for this article:

**Table S1** Site coordinates

**Table S2** Summary of effects of leaf economics traits on relative biomass, all sites

**Table S3** ANOVA table, effects of leaf economics traits on relative biomass

**Table S4** ANOVA table, effects of leaf economics traits on relative biomass, by site

**Table S5** ANOVA tables, effects of leaf economics GBLUPs on biomass GBLUPs

**Figure S1** Effects of genotype on leaf economics traits at putative QTL markers

**Figure S2** Effects of LMA and $N_{MASS}$ on relative biomass, by site

**Figure S3** Effects of $A_{MASS}$ and $N_{MASS}$ on relative biomass, by site

**Table S1** Site coordinates

| Site Code | Latitude | Longitude | City | State |
|---|---|---|---|---|
| KING | 27.55 | -97.88 | Kingsville | TX |
| PKLE | 30.38 | -97.73 | Austin | TX |
| CLMB | 38.9 | -92.22 | Columbia | MO |
| KBSM | 42.42 | -85.37 | Hickory Corners | MI |
| BRKG | 44.31 | -96.67 | Aurora | SD |

**Table S2** Table showing standardized effects of leaf economics traits on plant relative biomass across all sites. Estimate is parameter estimate from model; all quadratic term estimates are doubled

| **Parameter** | **Value** | **Std.Error** |
|---|---|---|
| Intercept | 0.988 | 0.02 |
| SiteCLMB | 0.004 | 0.017 |
| SiteKBSM | 0.05 | 0.018 |
| SiteKING | 0.005 | 0.054 |
| SitePKLE | -0.055 | 0.037 |
| LMA | 0.026 | 0.023 |
| $N_{MASS}$ | 0.015 | 0.02 |
| $A_{MASS}$ | -0.009 | 0.024 |
| SiteCLMB × LMA | 0.014 | 0.02 |
| SiteKBSM × LMA | 0.035 | 0.021 |
| SiteKING × LMA | 0.094 | 0.063 |
| SitePKLE × LMA | 0.063 | 0.043 |
| SiteCLMB × $N_{MASS}$ | -0.02 | 0.018 |
| SiteKBSM × $N_{MASS}$ | -0.04 | 0.018 |
| SiteKING × $N_{MASS}$ | -0.096 | 0.055 |
| SitePKLE × $N_{MASS}$ | -0.019 | 0.038 |
| SiteCLMB × $A_{MASS}$ | 0.048 | 0.021 |
| SiteKBSM × $A_{MASS}$ | 0.069 | 0.022 |
| SiteKING × $A_{MASS}$ | 0.137 | 0.065 |
| SitePKLE × $A_{MASS}$ | 0.026 | 0.044 |
| $LMA^2$ | -0.019 | 0.031 |
| $N_{MASS}^2$ | -0.043 | 0.031 |
| $A_{MASS}^2$ | 0 | 0.039 |
| LMA × $N_{MASS}$ | -0.05 | 0.026 |
| LMA × $A_{MASS}$ | -0.031 | 0.031 |
| $N_{MASS}$ × $A_{MASS}$ | 0.001 | 0.025 |
| SiteCLMB × LMA | 0.009 | 0.021 |
| SiteKBSM × LMA | 0.023 | 0.022 |
| SiteKING × LMA | 0.083 | 0.066 |

| | | |
|---|---|---|
| SitePKLE × LMA | 0.017 | 0.045 |
| SiteCLMB × $N_{MASS}$ | -0.033 | 0.018 |
| SiteKBSM × $N_{MASS}$ | -0.046 | 0.019 |
| SiteKING × $N_{MASS}$ | -0.111 | 0.056 |
| SitePKLE × $N_{MASS}$ | -0.024 | 0.038 |
| SiteCLMB × $A_{MASS}$ | 0.053 | 0.022 |
| SiteKBSM × $A_{MASS}$ | 0.075 | 0.022 |
| SiteKING × $A_{MASS}$ | 0.151 | 0.066 |
| SitePKLE × $A_{MASS}$ | 0.021 | 0.045 |
| SiteCLMB × $LMA^2$ | 0.017 | 0.026 |
| SiteKBSM × $LMA^2$ | 0.018 | 0.028 |
| SiteKING × $LMA^2$ | -0.018 | 0.082 |
| SitePKLE × $LMA^2$ | 0.101 | 0.056 |
| SiteCLMB × $N_{MASS}^2$ | 0.062 | 0.027 |
| SiteKBSM × $N_{MASS}^2$ | 0.001 | 0.027 |
| SiteKING × $N_{MASS}^2$ | 0.126 | 0.084 |
| SitePKLE × $N_{MASS}^2$ | 0.051 | 0.058 |
| SiteCLMB × $A_{MASS}^2$ | -0.03 | 0.036 |
| SiteKBSM × $A_{MASS}^2$ | -0.098 | 0.036 |
| SiteKING × $A_{MASS}^2$ | -0.137 | 0.109 |
| SitePKLE × $A_{MASS}^2$ | -0.08 | 0.075 |
| SiteCLMB × LMA × $N_{MASS}$ | 0.056 | 0.023 |
| SiteKBSM × LMA × $N_{MASS}$ | 0.024 | 0.024 |
| SiteKING × LMA × $N_{MASS}$ | 0.001 | 0.069 |
| SitePKLE × LMA × $N_{MASS}$ | -0.046 | 0.046 |
| SiteCLMB × LMA × $A_{MASS}$ | 0.01 | 0.027 |
| SiteKBSM × LMA × $A_{MASS}$ | -0.011 | 0.029 |
| SiteKING × LMA × $A_{MASS}$ | 0.025 | 0.085 |
| SitePKLE × LMA × $A_{MASS}$ | 0.053 | 0.061 |
| SiteCLMB × $N_{MASS}$ × $A_{MASS}$ | 0.063 | 0.025 |
| SiteKBSM × $N_{MASS}$ × $A_{MASS}$ | 0.07 | 0.023 |
| SiteKING × $N_{MASS}$ × $A_{MASS}$ | -0.05 | 0.07 |
| SitePKLE × $N_{MASS}$ × $A_{MASS}$ | -0.004 | 0.046 |

**Table S3** ANOVA table showing effects of leaf economics traits on plant relative biomass

| Parameter | df | F ratio | P |
|---|---|---|---|
| Intercept | 1, 2675 | 3378.73 | < 0.001 |
| Site | 4, 2675 | 3.57 | 0.007 |
| LMA | 1, 352 | 3.06 | 0.081 |
| $N_{MASS}$ | 1, 352 | 0 | 0.981 |
| $A_{MASS}$ | 1, 352 | 1.92 | 0.167 |
| Site × LMA | 4, 2675 | 0.97 | 0.421 |
| Site × $N_{MASS}$ | 4, 2675 | 1.11 | 0.352 |
| Site × $A_{MASS}$ | 4, 2675 | 3.43 | 0.008 |
| $LMA^2$ | 1, 346 | 1.87 | 0.172 |
| $N_{MASS}^2$ | 1, 346 | 0.18 | 0.667 |
| $A_{MASS}^2$ | 1, 346 | 0 | 0.995 |
| LMA × $N_{MASS}$ | 1, 346 | 6.01 | 0.015 |
| LMA × $A_{MASS}$ | 1, 346 | 0.81 | 0.368 |
| $N_{MASS}$ × $A_{MASS}$ | 1, 346 | 2.92 | 0.089 |
| Site × $LMA^2$ | 4, 2651 | 1.28 | 0.276 |
| Site × $N_{MASS}^2$ | 4, 2651 | 1.86 | 0.115 |
| Site × $A_{MASS}^2$ | 4, 2651 | 3.09 | 0.015 |
| Site × LMA × $N_{MASS}$ | 4, 2651 | 1.31 | 0.265 |
| Site × LMA × $A_{MASS}$ | 4, 2651 | 0.31 | 0.87 |
| Site × $N_{MASS}$ × $A_{MASS}$ | 4, 2651 | 3.41 | 0.009 |

**Table S4** ANOVA table showing effects of leaf economics traits on plant relative biomass, separately by site. Sites are ordered from north to south

| Site | Parameter | df | F ratio | P |
|---|---|---|---|---|
| **BRKG** | LMA | 1,352 | 1.2 | 0.274 |
| | $N_{MASS}$ | 1,352 | 0.53 | 0.468 |
| | $A_{MASS}$ | 1,352 | 0.15 | 0.701 |
| | $LMA^2$ | 1,346 | 0.38 | 0.536 |
| | $N_{MASS}^2$ | 1,346 | 1.94 | 0.165 |
| | $A_{MASS}^2$ | 1,346 | 0 | 0.991 |
| | LMA × $N_{MASS}$ | 1,346 | 3.6 | 0.059 |
| | LMA × $A_{MASS}$ | 1,346 | 0.99 | 0.321 |
| | $N_{MASS}$ × $A_{MASS}$ | 1,346 | 0 | 0.964 |
| **KBSM** | LMA | 1,352 | 6.13 | 0.014 |
| | $N_{MASS}$ | 1,352 | 1.36 | 0.244 |
| | $A_{MASS}$ | 1,352 | 5.95 | 0.015 |
| | $LMA^2$ | 1,346 | 0 | 0.988 |
| | $N_{MASS}^2$ | 1,346 | 1.71 | 0.192 |
| | $A_{MASS}^2$ | 1,346 | 5.57 | 0.019 |
| | LMA × $N_{MASS}$ | 1,346 | 0.92 | 0.339 |
| | LMA × $A_{MASS}$ | 1,346 | 1.53 | 0.217 |
| | $N_{MASS}$ × $A_{MASS}$ | 1,346 | 7.28 | 0.007 |

| | | | | |
|---|---|---|---|---|
| **CLMB** | LMA | 1,352 | 2.8 | 0.095 |
| | $N_{MASS}$ | 1,352 | 0.05 | 0.82 |
| | $A_{MASS}$ | 1,352 | 2.63 | 0.105 |
| | $LMA^2$ | 1,346 | 0 | 0.955 |
| | $N_{MASS}^2$ | 1,346 | 0.38 | 0.541 |
| | $A_{MASS}^2$ | 1,346 | 0.53 | 0.467 |
| | LMA × $N_{MASS}$ | 1,346 | 0.06 | 0.815 |
| | LMA × $A_{MASS}$ | 1,346 | 0.42 | 0.518 |
| | $N_{MASS}$ × $A_{MASS}$ | 1,346 | 5.7 | 0.018 |
| **PKLE** | LMA | 1,352 | 4.02 | 0.046 |
| | $N_{MASS}$ | 1,352 | 0.01 | 0.914 |
| | $A_{MASS}$ | 1,352 | 0.13 | 0.715 |
| | $LMA^2$ | 1,346 | 1.94 | 0.165 |
| | $N_{MASS}^2$ | 1,346 | 0.02 | 0.892 |
| | $A_{MASS}^2$ | 1,346 | 1.07 | 0.302 |
| | LMA × $N_{MASS}$ | 1,346 | 4.03 | 0.045 |
| | LMA × $A_{MASS}$ | 1,346 | 0.13 | 0.723 |
| | $N_{MASS}$ × $A_{MASS}$ | 1,346 | 0 | 0.945 |
| **KING** | LMA | 1,352 | 3.47 | 0.063 |
| | $N_{MASS}$ | 1,352 | 2.08 | 0.15 |
| | $A_{MASS}$ | 1,352 | 3.78 | 0.053 |
| | $LMA^2$ | 1,346 | 0.2 | 0.657 |
| | $N_{MASS}^2$ | 1,346 | 0.93 | 0.335 |
| | $A_{MASS}^2$ | 1,346 | 1.53 | 0.217 |
| | LMA × $N_{MASS}$ | 1,346 | 0.49 | 0.486 |
| | LMA × $A_{MASS}$ | 1,346 | 0 | 0.948 |
| | $N_{MASS}$ × $A_{MASS}$ | 1,346 | 0.47 | 0.492 |

**Table S5** ANOVA tables showing effects of leaf economics genomic linear unbiased predictors (GBLUPs) on plant biomass GBLUPs. GBLUPs were calculated from random effects models that included a random effect of kinship and a site-level random effect for biomass

| **Parameter** | **df** | **F ratio** | **P** |
|---|---|---|---|
| Intercept | 1, 346 | 0.16 | 0.690 |
| LMA | 1, 346 | 12.51 | <0.001 |
| $N_{MASS}$ | 1, 346 | 0.87 | 0.351 |
| LMA × $N_{MASS}$ | 1, 346 | 3.07 | 0.080 |

| **Parameter** | **df** | **F ratio** | **P** |
|---|---|---|---|
| Intercept | 1, 346 | 0.16 | 0.691 |
| $N_{MASS}$ | 1, 346 | 3.33 | 0.069 |
| $A_{MASS}$ | 1, 346 | 5.30 | 0.021 |
| $N_{MASS}$ × $A_{MASS}$ | 1, 346 | 4.66 | 0.032 |

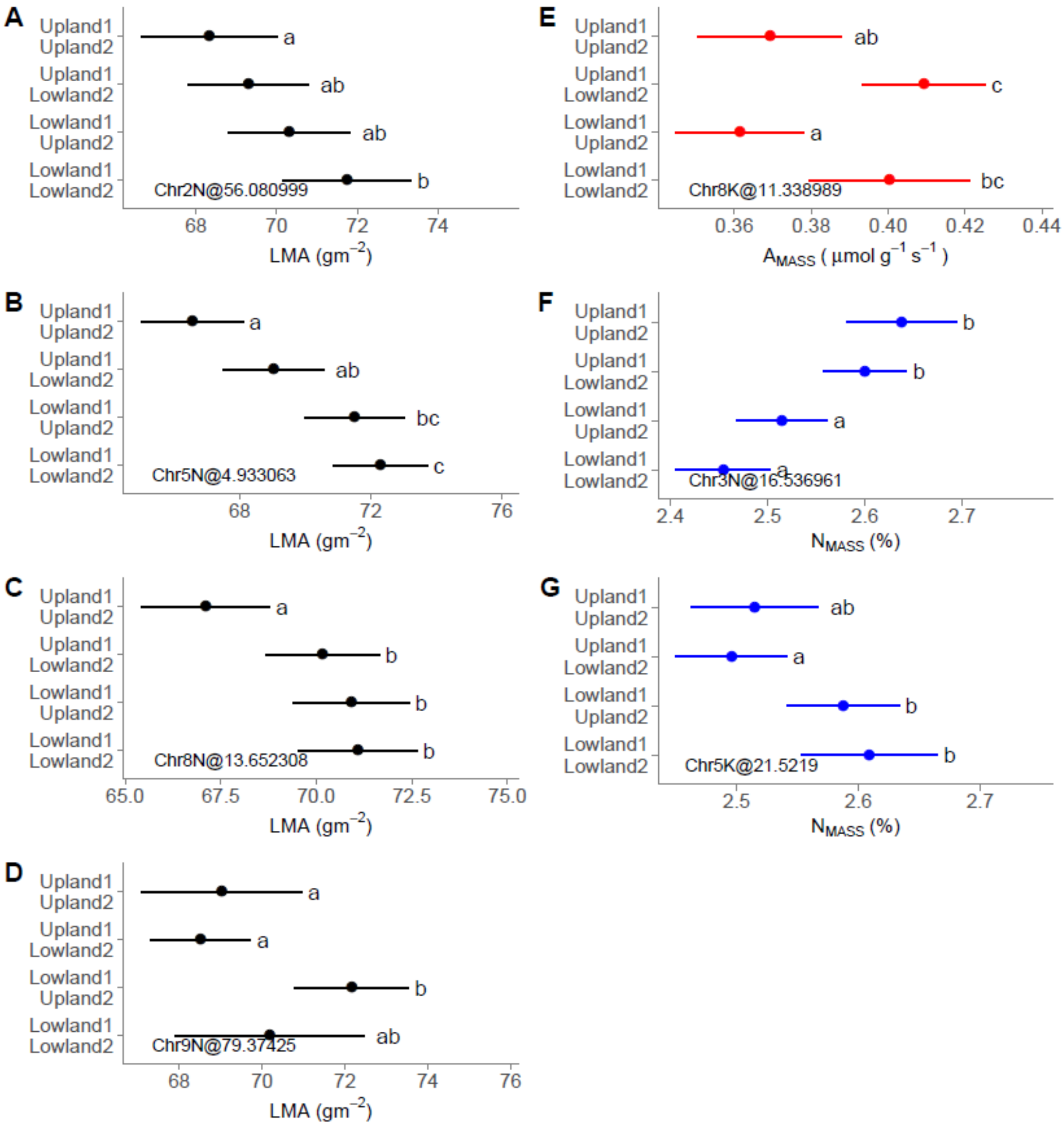


**Figure S1** Effects of genotype on leaf economics traits at putative QTL markers. Shared letters indicate no significant difference between groups

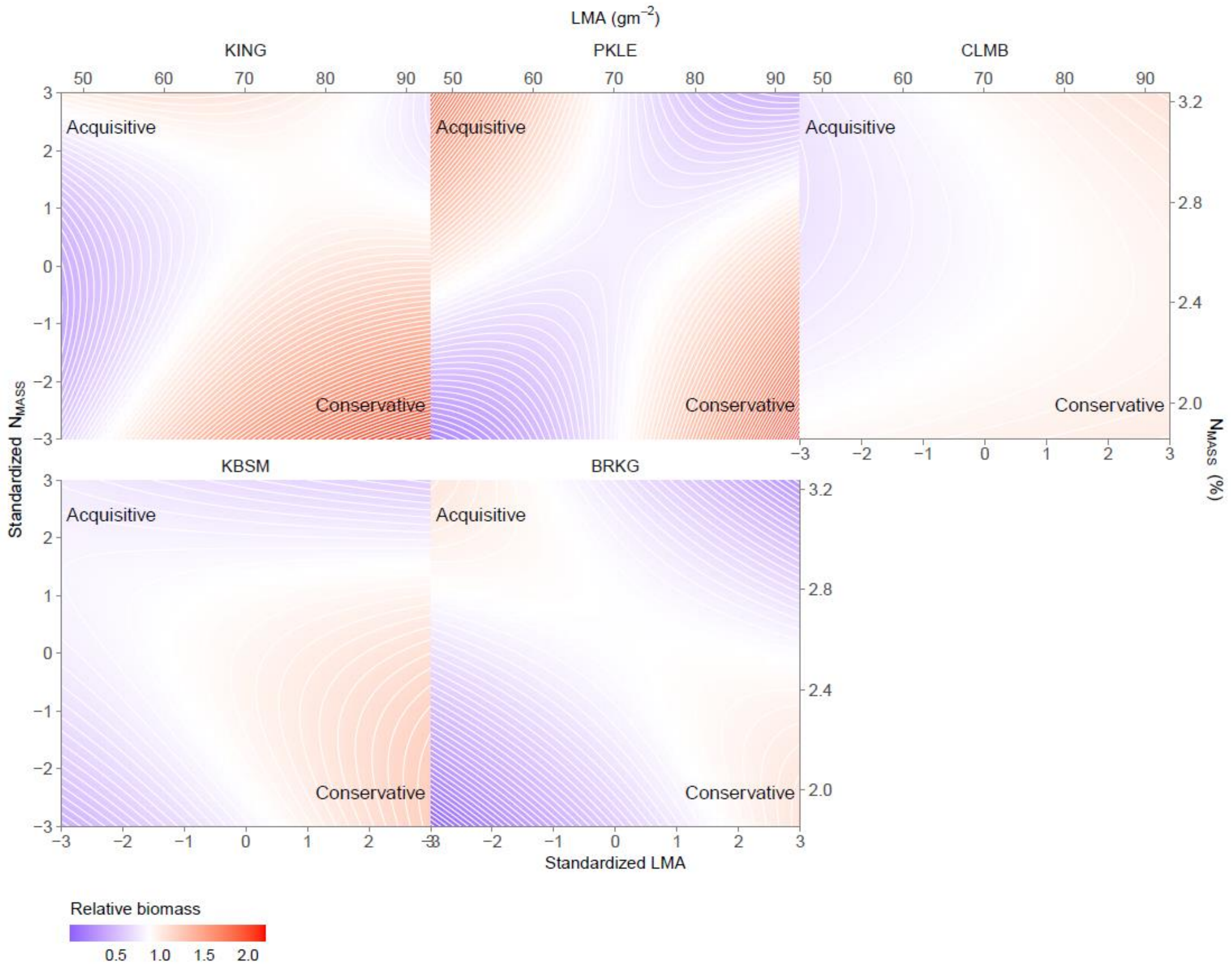


**Fig S2** Effects of standardized LMA and $N_{MASS}$ (transformed such that across all plants, mean = 0 and sd = 1) on relative biomass while controlling for $A_{MASS}$, separately by site. Model-derived quadratic parameter estimates are doubled

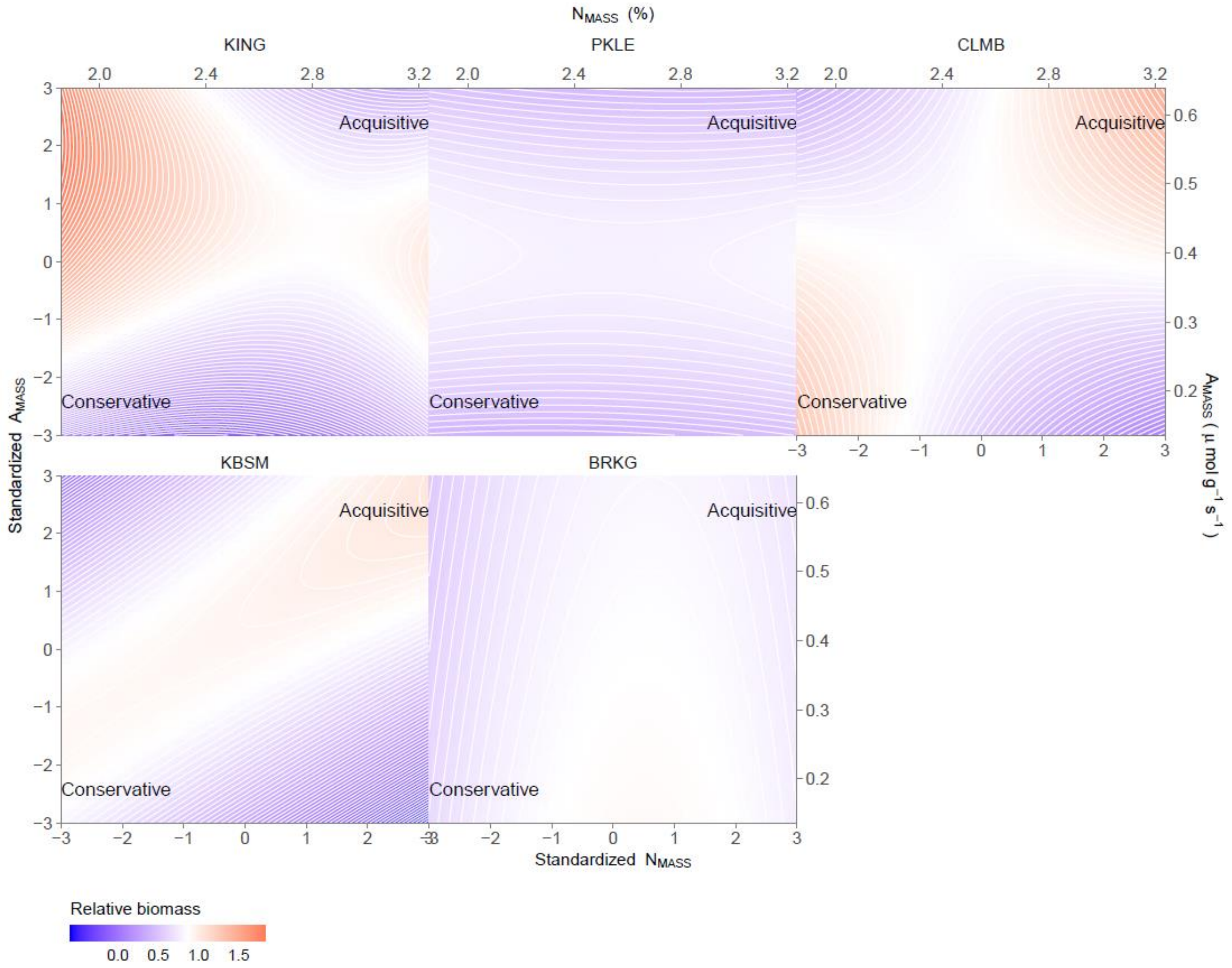


**Fig S3** Effects of standardized $A_{MASS}$ and $N_{MASS}$ (transformed such that across all plants, mean = 0 and sd = 1) on relative biomass while controlling for LMA, separately by site. Model-derived quadratic parameter estimates are doubled
